# Genome-wide comparative analysis reveals human-mouse regulatory landscape and evolution

**DOI:** 10.1101/010926

**Authors:** Olgert Denas, Richard Sandstrom, Yong Cheng, Kathryn Beal, Javier Herrero, Ross C. Hardison, James Taylor

## Abstract

**Background:** Because species-specific gene expression is driven by species-specific regulation, understanding the relationship between sequence and function of the regulatory regions in different species will help elucidate how differences among species arise. Despite active experimental and computational research, the relationships among sequence, conservation, and function are still poorly understood.

**Results:** We compared transcription factor occupied segments (TFos) for 116 human and 35 mouse TFs in 546 human and 125 mouse cell types and tissues from the Human and the Mouse ENCODE projects. We based the map between human and mouse TFos on a one-to-one nucleotide cross-species mapper, bnMapper, that utilizes whole genome alignments (WGA).

Our analysis shows that TFos are under evolutionary constraint, but a substantial portion (25.1% of mouse and 25.85% of human on average) of the TFos does not have a homologous sequence on the other species; this portion varies among cell types and TFs. Furthermore, 47.67% and 57.01% of the homologous TFos sequence shows binding activity on the other species for human and mouse respectively. However, 79.87% and 69.22% is repurposed such that it binds the same TF in different cells or different TFs in the same cells. Remarkably, within the set of TFos not showing conservation of occupancy, the corresponding genome regions in the other species are preferred locations of novel TFos. These events suggest that a substantial amount of functional regulatory sequences is exapted from other biochemically active genomic material.

Despite substantial repurposing of TFos, we did not find substantial changes in their predicted target genes, suggesting that CRMs buffer evolutionary events allowing little or no change in the TF – target gene associations. Thus, the small portion of TFos with strictly conserved occupancy underestimates the degree of conservation of regulatory interactions.

**Conclusion:** We mapped regulatory sequences from an extensive number of TFs and cell types between human and mouse. A comparative analysis of this correspondence unveiled the extent of the shared regulatory sequence across TFs and cell types under study. Importantly, a large part of the shared regulatory sequence repurposed on the other species. This sequence, fueled by turnover events, provides a strong case for exaptation in regulatory elements.

## Background

Most eukaryotic gene regulation occurs at the level of transcription (Derman et al. 1981) (Roop et al. 1978). This form of regulation involves the interaction of transcription factors (TFs) with element and function specific DNA sequences, referred to as *cis*-regulatory modules (CRMs; reviewed by (Maston et al. 2006)). Their modular organization allows for elaborate regulatory mechanisms and fine control of gene expression (Davidson 2006).

Evolutionary changes in CRMs have a profound effect on species divergence. Many studies suggest that species specific CRMs are the defining factor for species identity (Meader et al. 2010) (Taft et al. 2007) (Britten and Davidson 1969). Differences in human and chimpanzee, for example, are almost completely due to changes in functional noncoding sequence (King and Wilson 1975). Efforts to locate the “human gene” have only revealed differences in a small number of genes (Hughes and Yeager 1998), (Swanson et al. 2001), (Enard et al. 2002). Moreover, comparisons of organisms at the extremes of eukaryotes show that the genes encoding TFs and signaling components (e.g. for temporal/spatial gene expression patterns) are largely conserved (Davidson 2006). Taken together, this evidence suggests a hierarchical organization of regulatory networks. Modules at the top, performing essential upstream functions, span large evolutionary distances virtually unchanged, while lower level modules, involved in peripheral sub-networks, show a higher level of adaptation (Davidson and Erwin 2006). Under this model, part of the regulatory material must be under purifying selection and thus conserved between any two species of sufficiently small evolutionary divergence.

Evidence of conservation of regulatory sequence among species has inspired a series of computational methods. Some methods use machine learning and phylogenetic profiling to discover CRMs (reviewed by (Hardison and Taylor 2012)), others use comparative analysis to identify genomic material under purifying evolutionary constraint as a representative of the functional genome (reviewed by (Ponting and Hardison 2011)).

At the same time a number of experimental studies suggest that while sequence might encode enough information to drive TF binding (Wilson et al. 2008), the way this information is encoded is not trivial – similar sequence does not necessarily translate in similar function and vice versa. For example, regulatory elements have been found to tolerate sequence rearrangements (Hare et al. 2008) or even be under positive selection while maintaining the downstream regulatory machinery unchanged (Pheasant and Mattick 2007) (Bustamante et al. 2005). Recently, it was found that GATA1 changes its motif preferences during cell differentiation (May et al. 2013), serving as an example of TFs having multiple preferred motif sequences (Johnson et al. 2007) while maintaining a regulatory function.

Both computational and experimental approaches have provided valuable insight into the regulatory portion of the genome, but they have limitations. Computational methods are biased toward well-annotated and evolutionarily conserved genomic regions, indeed comparative analyses based on evolutionary conservation alone ignore species-specific functional elements. Experimental approaches based on direct genome wide measurements of TFos are a powerful resource for the identification and analysis of TF binding sites. However, the number of cell types and TFs assessed so far has often been limited.

With some studies pointing at conservation, others at divergence, and others yet at turnover of motifs and the importance of occupancy, the level if constraint on the CRMs is still an open question. This apparently contradicting evidence can be reconciled by considering conservation as specific to the TF or the cell type.

In this study, we combined several types of function-associated datasets from a large number of TFs from a wide variety of tissues and cell lines in both mouse and human. Our main data source is ChIP-Seq experiments performed by the mouse and human ENCODE projects. These data give evidence on the locations where TF have come close enough to the DNA to cross-link in cells. The question remains whether these locations, termed TFos (Hardison and Taylor 2012), could represent direct binding to a specific motif in the DNA or co-association with another TF. However, there is evidence that the TFos are active in assays for regulatory function at a far higher rate than non-occupied DNA segments, or DNA segments predicted as regulatory based on sequence motifs or conservation making them likely to have regulatory function. We also used DNase I Hypersensitive Sites (DHS) generated by the Human (The ENCODE Project Consortium et al. 2012) and the Mouse ENCODE (The Mouse ENCODE Consortium et al. 2014) projects. DHS regions are markers of regulatory DNA and have underpinned the discovery of all classes of *cis*-regulatory elements including enhancers, promoters, insulators, silencers and locus control regions. We conduced a comparative analysis, by integrating TFos with human-mouse WGA, gene annotations, and TF-gene associations. We have also compiled the data and annotations derived from this study into a database to serve as a resource in exploring the relationship between sequence evolution and function of regulatory elements **(Figure S1)**.

## Results

### An alignment based map for human-mouse TFos

Differences between present day genomes are the result of a series of evolutionary events originating on their most recent common ancestor (Kimura 1968). Many of these events can be explained under probability models and represented in the form of WGA (See (Ureta-Vidal et al. 2003) for a review). Sequence that has undergone a moderate number of evolutionary events will appear aligned, while highly divergent regions will usually be left un-aligned.

**Figure 1.**
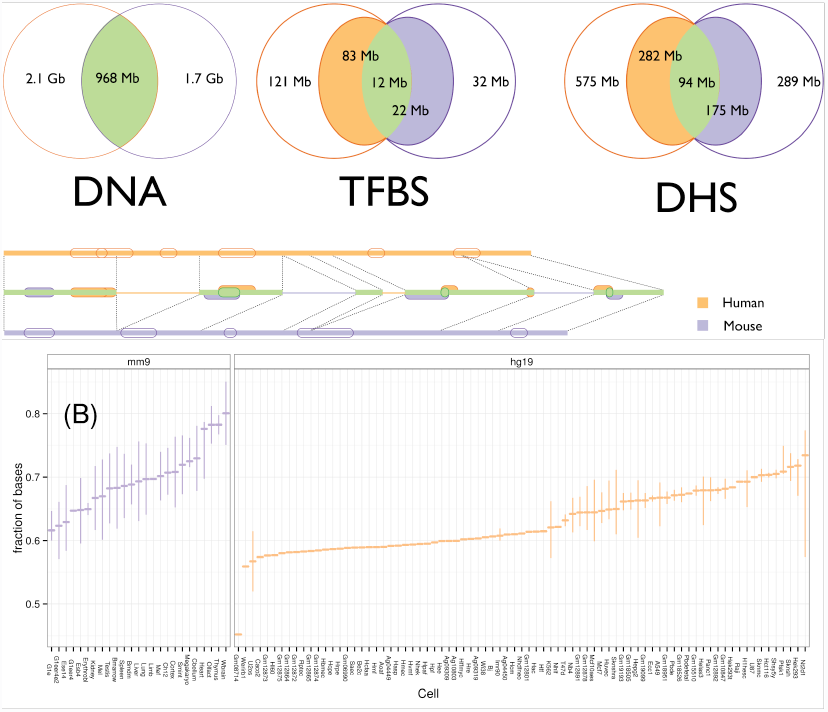
We built a one to one map between the human and mouse genomes from their WGA.

(A) The track diagram shows TFos (unfilled features on outer tracks) and how they map to the other species (middle track). The thick part of the middle track is the mappable DNA. The features on the middle track can be mapped on the other species and are: SeqCons if not overlapping features from the other species (respective color), or FuncCons or FunctActive otherwise (green). The Venn diagrams use the same color code to show the amount (rounded to the closest Mb) of mappable material from one species to the other for DNA sequence, TFos, and DHS. For example, human TFos cover 121 Mb of the human genome. When mapped to mouse, these TFos cover 83 Mb of the mouse genome and 12 Mb of the sequence covered also by mouse TFos. The diagram for DHS is labeled in a similar fashion.
(B) The distribution of mappable TFos nucleotides across cell types. The box-plot for each cell type summarizes the distribution of values for the fraction of nucleotides covered by TFos’s that can be mapped on the other species.

We used WGA to obtain a consistent map for TFos across human and mouse (Figure 1(A)). These alignments provide long, inferred homologous regions in the form of chains of gapless blocks. They can account for inversions, translocations, duplications, large-scale deletions, and overlapping independent deletions. We considered chained blastz (Schwartz et al. 2003) alignments available from the UCSC browser (Dreszer et al. 2012) and the 12-way mammalian WGA from the EPO pipeline (Paten et al. 2008) available from Ensembl version 65 (Flicek et al. 2013). To choose the most appropriate alignment for our mappings we used several criteria: symmetry, coverage, feature enrichment, and other method specific properties.

Symmetry is important for unambiguously mapping features from one genome to the other and back. The EPO based map is inherently symmetric; we only need to extract human-mouse alignments. We remove segmental duplications based on criteria such as the score or the length of the alignment to obtain an unambiguous map. UCSC alignments, on the other hand, are based on blastz pairwise alignments, which are not symmetric – a human-mouse alignment is different from a mouse-human alignment in general. However, it is possible to circumvent the problem by using a netting procedure and chaining again only the first layer in the resulting net. This corresponds to heuristically cleaning overlapping chains based on their score. The resulting reciprocal map is identical to the one derived from the mouse-human alignments at the cost of losing around 10% sequence coverage on both species (Kent et al. 2003).

UCSC alignments align a slightly larger fraction of the mouse genome (31% vs. 28% for UCSC and EPO respectively) to human. However, EPO alignments can assign a substantially higher amount of inserted mouse sequence (28% vs. 1.9%). In total 65.7% of the mouse genome remains unmapped by UCSC alignments and 42.5% by EPO alignments.

Next, we computed the number of features that each alignment could map on the other species for all available cell types in human and mouse. We found that UCSC alignments mapped more features on the other species **(Figure S2 and Table S4, S5, S6)**.

Based on the above considerations, we adopted the reciprocal UCSC alignments. The alignments and the comparative pipeline described here were adopted by the Mouse Encode Consortium for the cross-species mapping of TFose.

### Function and sequence conservation of TFos

We processed TFos generated by ChIP-Seq for 206 human and 55 mouse cell lines and tissues (The Mouse ENCODE Consortium et al. 2014), with a variable number of factors for each cell or tissue (from 1 to 109 for human and from 1 to 38 for mouse). The data were generated by the Human and the Mouse ENCODE projects. The ChIP-Seq experiments on mouse were conducted following the human ENCODE guidelines (Landt et al. 2012). Data processing, including Irreproducible Discovery Rate (IDR) analysis, was done using a uniform data processing pipeline for both datasets.

Elements recovered by the ChIP-Seq experiments and the subsequent peak-calling pipeline were subjected to thresholds for False Discovery Rate at 1% and IDR at 2%. We then filtered these sets to retain only those TFos showing DHS enrichment, thereby increasing our confidence in the functional role of the filtered elements. Roughly 41.63% of ChIP-Seq peaks were filtered out from each species by DHS filtering. The resulting data contained 5330864 and 727680 elements covering 121.08Mb and 31.6Mb of the genome for human and mouse respectively **(Table S3)**. The higher human coverage is due to the larger number of assays available in human.

We asked whether the selected putative regulatory material is significantly conserved. At the sequence level, we find that the fraction of TFos intersecting homologous regions is higher than one would expect by chance (Binomial test 0.99% CI: (0.7218, 0.7228) and 0.99% CI: (0.6908, 0.6936) with expected values of 0.3129 and 0.3648 for human and mouse respectively). At the TF-binding activity level (i.e., having TF occupancy by any TF), mapped human and mouse TFos overlapped TFos on the other species more often than expected by chance (Binomial test 0.99% CI: (0.5358, 0.5371) and 0.99% CI: (0.7883, 0.7913) with expected values 0.0231 and 0.0857 for human and mouse respectively). Significant conservation at the TF-binding activity level continues to hold largely for individual TFs-cell pairs **(Figures S3-S4)**. These results indicate that the most of these human and mouse regulatory regions have been under selective pressure both at the sequence and functional level.

Despite the selective pressure on the TFos, the data suggest extensive evolution of the regulatory material between human and mouse. On average 74.2% and 74.9% of TFos can be mapped on the other species resulting in a 36.1% and 31.4% of regulatory sequence coverage that is lineage-specific for human and mouse respectively (Figure 1(B) and **S5-12**). Importantly, we found that the extent to which the remaining regulatory material is conserved varies substantially between species and among cell types and TFs. Some previous studies on smaller numbers of TFs and cell types have emphasized conservation of TFos (Kim et al. 2007), while others find extensive regulatory sequence evolution (Odom et al. 2007). Thus our analysis of data for this much larger number of TFs and cell types gives a broader view of the extent to which TFos are conserved at the sequence and TF-binding activity level.

### Binding signal differences between functionally and sequence conserved TFos

We focus our TFos comparative analysis on two Tier 1 ENCODE cell lines and their mouse analogs as chosen by the Mouse ENCODE consortium. The data consist of 17 TFs from human chronic myelogenous leukemia cell line (K562, (Lozzio and Lozzio 1975)) and mouse erythroleukemia cell line (MEL, (The Mouse ENCODE Consortium et al. 2012)); and 15 TFs from lymphoblast cell lines (GM12878 vs. CH12) **(Table S2)**. In total we have thus selected 442527 and 215251 TFos covering 33.61Mb and 18.77Mb on human and mouse respectively. We mapped TFos of a species to homologous locations in the other and asked what function do those sites show.

**Figure 2.**
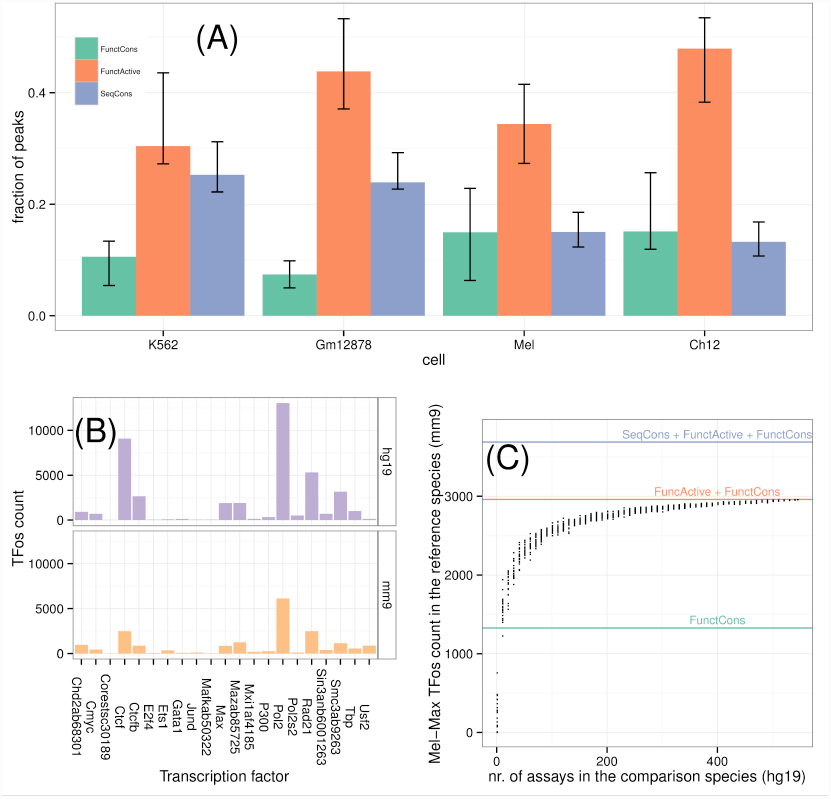
Conservation and re-use of TFos

(A) The distribution of SeqCons, FunctActive, and FunctCons regulatory elements summarized for all human mouse analogous cell lines **(Table S2)**. In both cell lines, a small fraction of mappable regulatory elements is FunctCons (33% and 15% cell average for human and mouse respectively). A larger fraction (46% and 50% cell average for human and mouse respectively) is FunctActive (plots for each cell are in **Figure S15, S16**).
(B) Loss and gain of TF binding sites. We used TFos in our data to discover TF binding sites (TFBS) on both species. We mapped human (respectively mouse) TFBS to mouse (respectively human) and computed their distance to the closest mouse (respectively human) TFBS of the same TF on an analogous cell. Among these TFBS those with a positive distance but less than 150bp contribute to the corresponding bar in the human (respectively mouse) subplot. In other words, each bar is the count of TFBS that were lost and gained within a 150bp window of the original site.
(C) Reuse of FunctActive TFos between different cells and factors. A point on the scatterplot indicates the number of TFos of the corresponding assay in the reference species that can be classified as FunctCons or FunctActive when considering a fixed size set of randomly chosen assays in the comparison species. We performed multiple computations for each chosen size of such sets. Lines indicate the accumulated number of FunctCons, FunctActive, and Seqcons TFos. In this figure, about 93% of the (Mel, Max) assay is covered by TFos from just 35% of the query assays (plots for other cells/TFs in **Figures S17, S18**).

The homologous site of a TFos can be: (i) not occupied by TFs, in which case we call the TFos *SeqCons*; (ii) repurposed, thus active in another cell or bound by another factor in the same cell type, in which case we call the TFos *FunctActive*; or (iii) bound by the same factor on the same cell type, in which case the TFos is called *FunctCons*. SeqCons and FunctActive elements represent differences in human and mouse TF binding patterns due to TFos loss, gain, or both. Turnover occurs in the case of loss and gain of the same TFos at different, but nearby, positions. However, more complex differences can arise when the loss of one TFos is followed by compensatory gains of TFos of other TFs. For the pairs of analogous cell types in human and mouse, the largest proportion of TFos (considered together) are FunctActive, while the relative proportions of FunctCons and SeqCons vary between cell types and species (Figure 2(A)).

Conservation of occupancy in the other species was associated with peak signal strength for several cell type-TF pairs. For example, binding signal on FunctCons or FunctActive elements was higher than that on SeqCons elements for 59% and 50% of cell type-TF combinations for human and mouse respectively (with a Bonferroni-corrected error rate of 1%; **Figures S13, S14**). Overall, FunctCons peak signal is on average 1.3 and 1.9 times larger than SeqCons peak signal on human and mouse respectively **(Table S1)**.

### TF binding site turnover and TFos repurposing

If SeqCons and FunctActive elements are the result of turnover of TF binding sites (TFBS) in human and mouse, then we should observe TFBS close to regions orthologous to TFBS on the other species. We discovered TFBS using motif discovery and motif matching on TFos for both species and analyzed their alignment. We found that for several TFs, SeqCons TFBS map within 150bp of a TFBS on the other species. On average, 51% and 48% of SeqCons TFBS have been subject to turnover in a 150bp neighborhood for human and mouse respectively (Figure 2(B)).

The large number of FunctActive elements (Figures 2(A) and **S15, S16**) suggests recycling of TFos among cells and TFs. We examined this more carefully with a novel approach, based on the intuition that extensive recycling would allow any set of TFos in a comparison species to identify the occurrence of specific TFos in the reference species (after cross-species mapping). To conduct this analysis, we sampled (multiple times) mapped TFos from *k* assays from the comparison species, took the union of these TFos, and computed the number of TFos a given assay (in the reference species) shares with this union. The results strongly support extensive recycling of TFos. Specifically, we find that about 93% of the (Mel, Max) assay is covered by TFos from just 35% of the query assays. As *k* increases, coverage saturates (Figures 2(C) and **S17, S18**). This ability of different assays in the comparison species to capture TFos in the reference species is substantially different than what is obtained by simulating non-associated assays (**Figure S19**).

We tested the hypothesis that new TFos tend to arise over existing TFos – active in other cells or occupied by other factors. Under the null hypothesis, the fraction of FunctActive elements with respect to the non-FunctCons elements should not be significantly higher than the fraction of the genome that can turn into a FunctActive TFos, should a new TFos occur (See Materials and Methods for details). We found that fraction to be higher than expected (One sided Binomial test 99% CI: (0.377, 1.000) and (0.477, 1.000) with expected values 0.010 and 0.028 for human and mouse respectively), suggesting that existing CRMs are sites where new TFBS are likely to arise.

**Figure 3.**
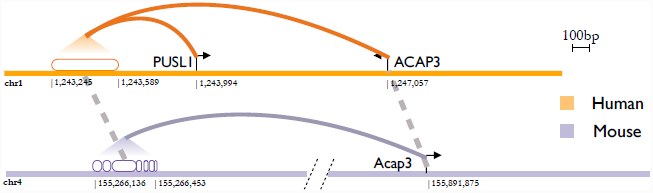
Conservation of presumptive gene targets for a repurposed TFos We combined cross-species gene-gene and TFos-TFos associations and gene-TFos associations to determine whether the sets of target genes of FunctCons and FunctActive TFos differ significantly. Gene-gene association data are based on a set of orthologous genes between human and mouse produced by the mouse ENCODE consortium, gene-TFos data are based on synchronized DHS activity during gene transcription (Thurman et al. 2012), and TFos-TFos associations are based on our cross-species map. In the figure, we show a human TFos of Mxi on K562 (empty oval) associated with PUSL1 and ACAP3. This TFos is FunctActive, since its analogous location (broken oval linked by the dashed line with spaces between ovals indicating insertions in mouse) in mouse is bound by other TFs on other cell types (not shown). However, its analogous site in mouse is linked to gene Acap3. Incidentally, ACAP3 and Acap3 are orthologous and in our gene-gene association set. The human TFos and its analogous site in mouse bind different TFs and are active in different cell types, but they share a target gene.

Turnover events or the appearance of novel TFBS lead to compositional changes in CRMs. However, these changes do not always reflect downstream in, for example, the set of target genes. Following (Thurman et al. 2012) we extracted a set of TSS-enhancer connections based on synchronized DHS activity during the transcription of a gene. Restricting on 15,736 human-mouse orthologous genes (The Mouse ENCODE Consortium et al. 2014), we were thus able to define putative target genes for 1928211 and 204758 TFos for human and mouse respectively. We observe that, similarly to FunctCons TFos, about 39% and 43% of FunctActive TFos retained at least one putative target gene for human and mouse respectively. Furthermore, the amount at which the set of target genes is conserved across species is not significantly different between the two classes of TFos (Kolmogorov-Smirnov test. p-values: < 2.2e-16 and < 2.2e-16 for human and mouse respectively). The example of the candidate enhancer linked to the *ACAP3* gene in human illustrates this situation (Figure 3). The DNA segment in mouse that is orthologous to the human enhancer has been repurposed such that it binds different TFs, but the mouse *Acap3* gene is still the presumptive target.

## Discussion

We reported on the construction and use of a map of functional elements between human and mouse based on WGA. This map is consistent and symmetrical in that it provides a one to one correspondence between genome elements in both directions. Furthermore, it constitutes an improvement over aligning TFos directly on the other species guided by gene orthology; this approach is likely to deteriorate with the distance of an element from the nearest gene with an ortholog on the other species.

Using this mapping approach, we were able to recover, on average, 75% and 73% of TFos in the other species, of which 63% and 78% are bound by any TFs and 13% and 25% by the same TF for human and mouse respectively. The rest of the regulatory material is species-specific either at the sequence or functional level and is likely to account for the phenotypic differences between human and mouse (King and Wilson 1975) (Davidson and Erwin 2006). However, some of the species-specific material retains target genes suggesting that CRMs buffer changes toward downstream regulation (Weirauch and Hughes 2010).

One potential difficulty, which affects this analysis, is the variation that can be introduced by technical factors such as signal thresholding or by differences in the environment of the cells being compared. By using data processed for reproducibility under the stringent standards developed by the Human and Mouse ENCODE projects, restricting to regions overlapping DHS for greater confidence, considering the TF activity on analogous cell types, and employing statistical controls we hope to have ameliorated these issues.

Previous studies have mapped the binding patterns of TFs between species, including mouse and human, by focusing on a single cell type and a few TFs (e.g., (Odom et al. 2007)). While these important studies have revealed substantial divergence in the binding patterns of TFs between species, they do not explore FunctActive elements or related dynamics such as repurposing. One important conclusion in our study is that the evolution of regulatory sequences varies considerably among TFs and cell types, and furthermore we document which TFos fall into the various evolutionary categories.

The traffic between noncoding functional and nonfunctional DNA in a genome is two-way. On the one hand, loss of a TFos can increase fitness, as dramatically illustrated by changes in the regulation of *pitx1* in three-spined stickleback that lead to advantageous anatomical changes in a new environment. On the other hand, new functional noncoding sequence can arise from nonfunctional sequence through turnover (Wilson and Odom 2009). The differences between the sizes of FunctActive and FunctCons pools suggests another source of functional noncoding material traffic, characterized by the exaptation of TFos into functional noncoding material with novel TF or cell type activity (Gould and Vrba 1982). With a partial catalog of regulatory elements it is hard to distinguish the newly created functional material from the exapted one. However with ever-increasing numbers of experimental assays the picture should become more clear.

## Methods

### Data processing

#### ChIP-Seq data

We download all of the ChIP-Seq data generated by the Human and the Mouse ENCODE from the ENCODE DCC. The pipeline that filters the original ENCODE peaks is available in the supplementary material.

#### Distal DHS-to-promoter connections

As described in (Thurman et al. 2012) many cell-selective enhancers become DHSs synchronously with the appearance of hypersensitivity at the promoter of their target gene. This has been used to infer a genome-wide DHS/enhancer-promoter connection set. Using a conservative list of orthologous genes (The Mouse ENCODE Consortium et al. 2014) we inferred a list of correspondences between TFos regulating human-mouse orthologous genes.

#### EPO and UCSC based maps

We considered the EPO 12-way mammalian whole genome alignments from Ensembl project version 65 (Flicek et al. 2013) and the UCSC human mouse chain and net alignments from the UCSC (Dreszer et al. 2012) genome browser to generate the reciprocal maps. Details and programs for processing of the alignments are available at https://bitbucket.org/bxlab/mapper_comparisons

#### Mapping strategy

We built and used a one-to-one nucleotide mapper (bnMapper) to map elements between human and mouse. The mapping is bijective, so reverse application of the mapping to a mapped nucleotide, returns the original nucleotide. Elements that span matching blocks of different chains and elements that map to multiple chromosomes are filtered out. The mapper and a detailed analysis of performance between EPO and UCSC alignments are available at https://bitbucket.org/bxlab/mapper_comparisons. Details on the mapping strategy are on Section 1.1 of the supplementary material.

### Significance tests

For a given genomic region of coverage *C*, the number of n randomly positioned features over the genome (length *L*) would follow a *Binomial(n, C/L)*. For significance of TF conservation at the sequence level we set *L, C*, and *n*, to be the length of the genome, total TF coverage, and total number of peaks. A success event is an overlap of 1bp with the one to one mappable sequence. Similarly, for functional conservation, *L, C*, and *n*, are set to total one to one mappable sequence, coverage of mapped peaks, and number of mapped peaks overlapping with peaks on the target genome, respectively.

For the paired Wilcoxon tests we consider all TFs within a cell line that appear on both species. Ranks are computed from FunctCons to SeqCons ratios of corresponding TFs on each species.

To test that a new TFos is more likely to occur in an existing TFos, possibly occupied by another TFs or active in another cell, consider the following quantities with respect to a fixed genome:

- The number of non-FunctCons TFos in this genome not less than the number of new TFos in this genome. The inequality accounts for TFos deletions on the other genome. We write *nFCo* > = *nFCt*.
- Coverage of SeqCons of the other genome after being mapped in this genome and FA elements in this genome not less than regions in this genome that would become FA should a new TFos occur. The inequality accounts for those regions that would become FunctCons should a new TFos appear. We write *Mo* >= *Mt*.
- Length of this genome. We write *L*.
- The number of FunctActive Tfos in this genome. We write *FA*.

Here we are making the assumption that FunctCons elements have existed before human-mouse split and FunctActive are a result of a deletion or creation (or both) after the human-mouse split. Thus, the fraction *FA / nFCt* is that of new elements that occurred in existing TFos involving other TFs or cell types, whether the fraction *Mt / L* indicates the chance of a new TFos to become FunctActive should it occur randomly in the genome. Under the alternative hypothesis, *FA / nFCt > Mt / L*. However, because *Mt / L <= Mo / L* and *FA / nFCo <= FA / nFCt*, it suffices to show that *FA / nFCo >= Mo / L*. Notice that we observe all the quantities in the last expression.

### Data access

UCSC alignments can be downloaded from the UCSC website http://hgdownload.soe.ucsc.edu/goldenPath/hg19/vsMm9/

UCSC bijective (netted) alignments, the list of one-to-one orthologous genes, the list of DHS-gene associations, and the list of Mouse ChIP-Seq TFos and DHSs can be downloaded from the Mouse ENCODE web portal http://mouse.encodedcc.org/data and http://www.mouseencode.org/

The mapping software can be freely downloaded as part of the bx-python software library from https://bitbucket.org/james_taylor/bx-python/wiki/Home

Ensembl EPO 12-way alignments version 65 can be downloaded from the Ensembl (December 2011) at ftp://ftp.ensembl.org/pub/release-65/emf/ensembl-compara/epo_12_eutherian/ and http://www.ebi.ac.uk/∼kbeal/species_mapper/epo_547_hs_mm_12way_mammals_65.out.gz

The human ChIP-Seq TFos and DHSs can be downloaded from the Human ENCODE website at http://genome.ucsc.edu/ENCODE/

## Acknowledgments

Thanks to members of the Taylor and Hardison labs and the Mouse ENCODE consortium for their input and discussion. This project was supported by grants from the National Institutes of Health to RCH and JT, specifically American Recovery and Reinvestment Act (ARRA) funds through grant number RC2 HG005573 from the National Human Genome Research Institute,and grant number R01 DK065806 from the National Institute for Diabetes, Digestive, and Kidney Diseases. This work was also supported by the Wellcome Trust (grant number 095908) and the European Molecular Biology Laboratory.

## Authors’ Contributions

OD wrote the main text, performed the comparative and statistical analysis, and wrote the mapper tool. RS helped designing the mapper, carried out statistical analysis for choosing the best whole genome alignments, prepared the DHS data, and the gene – TFos link data. YC helped on the design of the mapper and performed analyses on the whole genome alignments. KB worked on the design of the mapper and the preparation of the whole genome alignments. JH worked on the design of the mapper and the preparation of the whole genome alignments. He helped design of the study and write the main text. RC wrote the main text and designed the study – JT wrote the main text, carried out statistical analyses, and designed the study.

